# The double-drift illusion biases the marmoset oculomotor system

**DOI:** 10.1101/2023.02.02.526908

**Authors:** Nicholas M. Dotson, Zachary W. Davis, Jared M. Salisbury, Stephanie E. Palmer, Patrick Cavanagh, John H. Reynolds

**Affiliations:** The Salk Institute; The Salk Institute for Biological Studies; University of Chicago; Glendon College, York University; Salk Institute for Biological Studies

## Abstract

The double-drift illusion has two unique characteristics: the error between the perceived and physical position of the stimulus grows over time, and saccades to the moving target land much closer to the physical than the perceived location. These results suggest that the perceptual and saccade targeting systems integrate visual information over different timescales. Functional imaging studies in humans have revealed several potential cortical areas of interest, including the prefrontal cortex. However, we currently lack an animal model to study the neural mechanisms of location perception that underlie the double-drift illusion. To fill this gap, we trained two marmoset monkeys to fixate and then saccade to the double drift stimulus. In line with human observers for radial double-drift trajectories, we find that saccade endpoints do show a significant bias that is, as it is in humans, smaller than that seen in perception. This bias is modulated by changes in the external and internal speeds of the stimulus. These results demonstrate that the saccade targeting system of the marmoset monkey is influenced by the double-drift illusion.

## Introduction

Perceptual illusions provide an opportunity to uncover fundamental mechanisms of sensory processing (Gregory, 1968; Eagleman, 2001). The double-drift visual illusion is one example that has recently been shown to dissociate the positional representations in the perceptual and oculomotor systems (Lisi & Cavanagh, 2015). The double-drift illusion is created using a Gabor patch with an internal motion that is orthogonal to the aperture motion. When viewed in the periphery the patch appears to move in a direction ~45° from the actual direction (Fig. 1). This results in a position offset between the physical and perceived location that accumulates over time (Lisi & Cavanagh, 2015; Liu et. al., 2019; ’t Hart et. al., 2022). The double-drift illusion also demonstrates an intriguing dissociation between perceptual and saccadic reports. In human subjects asked to saccade to a double-drift stimulus moving vertically to the right or left of fixation, there is little or no effect of the illusion on saccade landings (Lisi & Cavanagh, 2015). This suggests that the saccade targeting system may be integrating information over a shorter timescale than the perceptual system. However, for double-drift stimuli moving toward or away from the fovea, the saccades land near the physical location but are reliably offset in the direction of illusory motion (Lisi & Cavanagh, 2022). This link between saccade bias and perception for saccades to radial target paths offers an opportunity to utilize saccadic reports in non-human primates as a proxy for a perceptual report.

**Figure 1:**
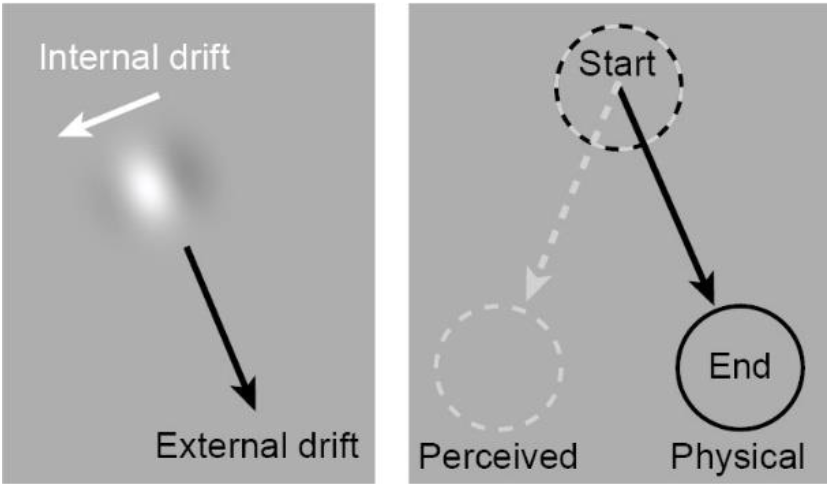
Illustration of the double-drift illusion. Fast internal motion combined with slower external drift produces a profound positional offset. In the case where the aperture is drifting to the lower right with an internal motion that is orthogonal and to the left (left panel), the stimulus will appear to be moving down and to the left (right panel).

Functional imaging studies in humans have revealed several potential cortical areas of interest, including the prefrontal cortex and visual motion processing areas of the visual system (Liu et. al., 2019; Steinberg et. al., 2022). However, we currently lack an animal model to study the neural mechanisms underlying the double drift illusion. Marmosets are an emerging model organism for visual neuroscience (Mitchell et. al., 2014; Solomon & Rosa, 2014; Davis et. al., 2020; D’Souza et. al., 2021), with several advantages over the commonly utilized macaque monkey, including a lissencephalic (smooth) cortex which facilitates array and laminar recordings in cortical areas that are buried within sulci in macaques (Johnston et. al., 2019; Selvanayagam et. al., 2019; Davis et. al., 2020; Davis et. al., 2022). This is particularly useful for studying cortical areas such as the frontal and visual cortex which have been shown in human subjects to be relevant for the double-drift illusion.

Here we test whether or not the marmoset oculomotor system is biased by the double-drift illusion. Marmosets were trained to perform the Double-Drift Saccade Task (DDST), which requires them to fixate and then wait a brief period of time before saccading to the double-drift stimulus. Similar to human observers, we find that their saccade endpoints show a bias that is modulated by changes in the external and internal velocity of the stimulus. These results demonstrate that the saccade targeting system of the marmoset monkey is biased by the illusion, indicating that they are a suitable model for studying the neural mechanisms of location perception that underlie the double-drift illusion.

## Methods

Two adult male marmoset monkeys (*Callithrix jacchus*) were used in this study. To stabilize the head and track the eye position a headpost was surgically implanted on each monkey. All surgical procedures were performed with the animal under general anesthesia in an aseptic environment in accordance with the recommendations in the Guide for the Care and Use of Laboratory Animals of the National Institutes of Health. All experimental methods were approved by the Institutional Animal Care and Use Committee (IACUC) of the Salk Institute for Biological Studies and conformed with NIH guidelines.

Marmosets were trained to freely enter a custom-made marmoset chair, which was then positioned 41 cm from a calibrated and gamma corrected LCD monitor (ASUS VG248QE; 100 Hz refresh rate; 75 cd/m_2_ background luminance). Eye position was measured with an IScan CCD infrared camera (500Hz sampling rate). MonkeyLogic was used to calibrate and record eye position (1 kHz sampling rate), along with stimulus presentation and behavioral control (Asaad & Eskandar, 2008; Hwang et. al., 2019).

Marmosets performed the Double-Drift Saccade Task (DDST). After fixation was acquired and held for 250 ms within a fixation window (2.25 degrees of visual angle (dva) radius) a drifting Gabor stimulus appeared at a random location 12 dva from the fixation point (Fig. 2). The stimulus traveled up to 6 dva over 1000ms at a slight angle (12 deg.) to the fixation point. The direction of internal motion was always orthogonal and inward with respect to the fixation point. On trials with no external motion, the stimulus remained in a fixed location (70% of the path length). After 500 ms the fixation point disappeared and the marmoset was allowed 500 ms to saccade to the stimulus. During the saccade period, the stimulus continued to travel along its path but was extinguished as soon as fixation was broken. To receive a reward, the eye position had to land in the saccade window (3.5 dva radius centered at the end of the stimulus path) within 100 ms after fixation was broken and remain within the window for 100 ms. These parameters ensured that 1) saccades were directed toward the stimulus without constraining the endpoints, 2) only one saccade was made to the stimulus, and 3) that the saccade landed within the saccade window rather than pass through it. If these criteria were met then a small reward (marshmallow fluff and water mixture (3g fluff : 1ml water)) was given using a syringe pump.

**Figure 2:**
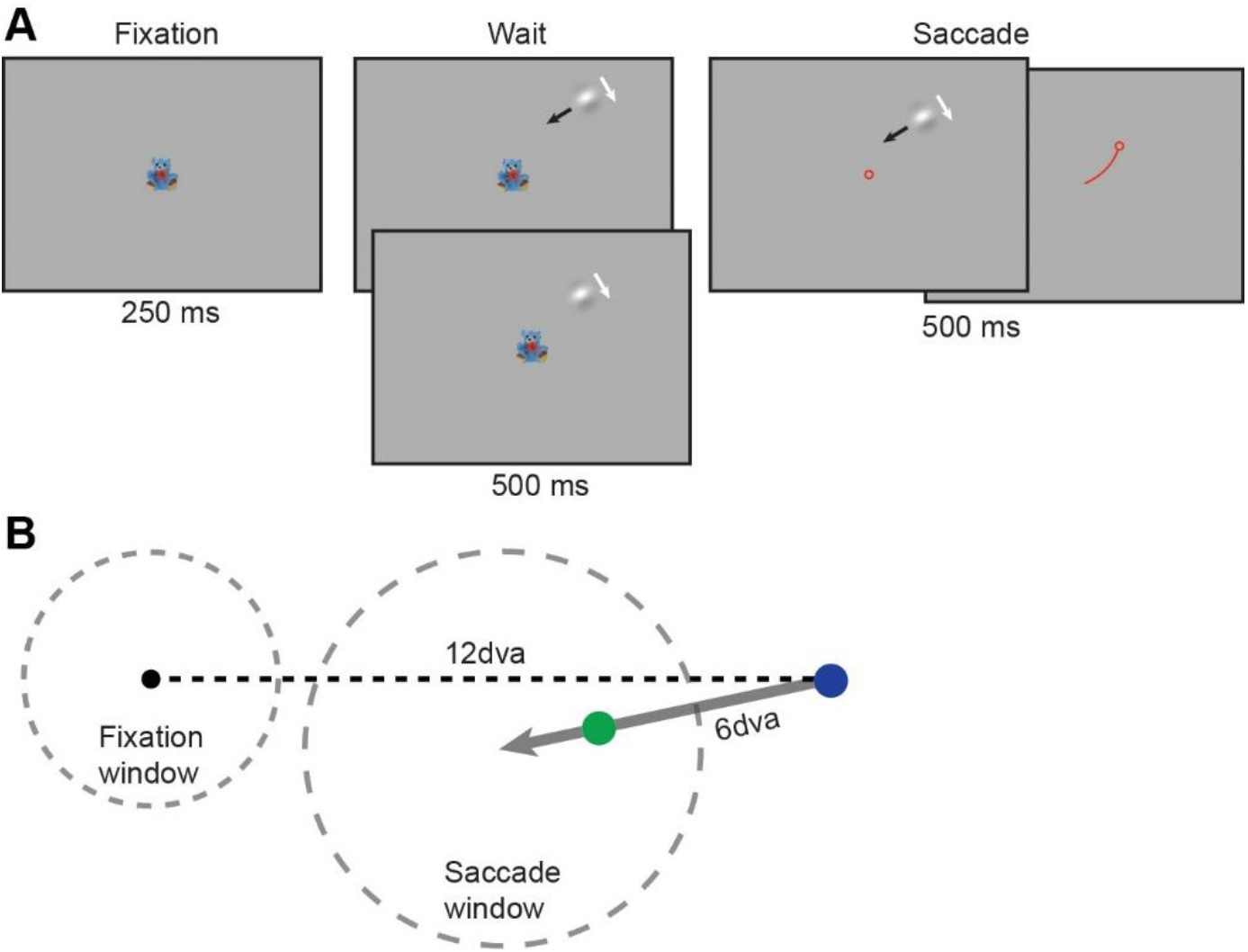
Illustration of the double drift saccade task (DDST). A) Marmosets were required to maintain fixation in the presence of the double drift stimulus and then saccade to the stimulus. The stimulus was extinguished at the onset of the saccade. The top and bottom panels in the Wait period indicate the conditions with and without external motion. In the Fixation, Wait, and Saccade panels the red circle indicates eye position. The Saccade panels illustrate that the stimulus remains on (left) until a saccade is made (right). In the rightmost Saccade panel, the red line indicates eye trajectory and the red circle indicates post-saccadic eye position. B) Diagram of the relative positions of the fixation point, stimulus path, and the fixation (2.25 dva radius) and saccade (3.5 dva radius) windows. The gray arrow indicates the stimulus path. The blue dot indicates the start position of the stimulus (non-zero external speed). The green dot indicates the position of the stimulus when the external speed is zero.

The Gabor patch (80% Michelson contrast) consisted of a sinusoid carrier with a spatial frequency of 0.4 cycle/dva and a Gaussian envelope with a standard deviation of 0.5 dva. The mean luminance matched the background luminance. In two of the conditions, the external speed was fixed at 6 dva/s, while the internal motion speed was either 4 dva/s or 8 dva/s. In the third condition, the external speed was 0 dva/s and the internal speed was 8 dva/s. Since the patch was not moving across the screen, the location was fixed at a point where the marmosets would typically saccade to the moving patch (70% of the full path length). Only one condition was used during a recording session. For all conditions, on 50% of the trials, the internal speed was 0.1dva/s (non-illusory). These trials, which do not result in an illusion in human observers, were used to control for any bias in each animal’s saccade endpoints, independent of the illusion.

Eye data noise was reduced offline by using a moving average (20 ms window). To determine the eye position during the Wait period, we combined eye position data across trials (500 sample points for each trial). Instantaneous eye velocity was calculated using the difference in eye position between samples separated by 10ms. The Matlab functions risetime.m and falltime.m were used to identify the start and stop times of the saccades based on the eye velocity. We calculated the angle bias as the angle between the saccade endpoint and the physical path of the stimulus relative to the start position of the stimulus. Saccade amplitudes were calculated as the distance between the center of fixation and the endpoint of the saccade. To account for any bias due to the configuration of the stimuli or eye calibration offsets, we subtracted the median angle of the non-illusory control trials from the illusory trials for each condition. A positive angle bias indicates that the saccade endpoint is on the side the internal motion is oriented toward (illusory direction).

## Results

We collected behavioral data from two marmoset monkeys performing the Double-Drift Saccade Task (DDST) under three different experimental conditions. For each condition, data are from four sessions, two from each monkey. The DDST requires the monkey to briefly hold fixation and then saccade to the stimulus (Fig. 2). During the Wait period, the monkey’s gaze was typically less than 1 dva from the center of the fixation window (median = 0.6 dva and 90% ≤ 1.25 dva). Based on results from human observers, we predicted that the saccades would be biased by the internal motion when parameters were in a range that caused strong perceptual effects (Lisi & Cavanagh, 2022; Massendari et al., 2018). In general, slow external motion combined with fast internal motion should lead to larger saccade biases than slow external motion combined with slow internal motion or just internal motion alone (Heller et. al., 2021). First, to determine if saccade endpoints are biased by the internal speed, we compared sessions with slow (4 dva/s) and fast (8 dva/s) internal speeds and a fixed external speed of 6 dva/s. Next, we kept the internal speed fixed at 8 dva/s and compared external speeds of 0 dva/s and 6 dva/s.

To determine if saccade endpoints are biased by the internal speed, we compared sessions with slow (4 dva/s) and fast (8 dva/s) internal speeds and a fixed external speed (6 dva/s) (Fig. 3A). In the slow (4 dva/s) condition, saccades were very slightly biased toward the direction of internal motion (median angle bias = 1.35 deg.; p = 0.03, Wilcoxon signed rank test). However, in the fast (8 dva/s) condition, saccade endpoints were strongly biased (median angle bias = 4.93 deg.; p = 1.1E-7, Wilcoxon signed rank test), and we found a large difference in the angle bias between the two conditions (p = 4.9E-4, Wilcoxon rank sum test).

**Figure 3:**
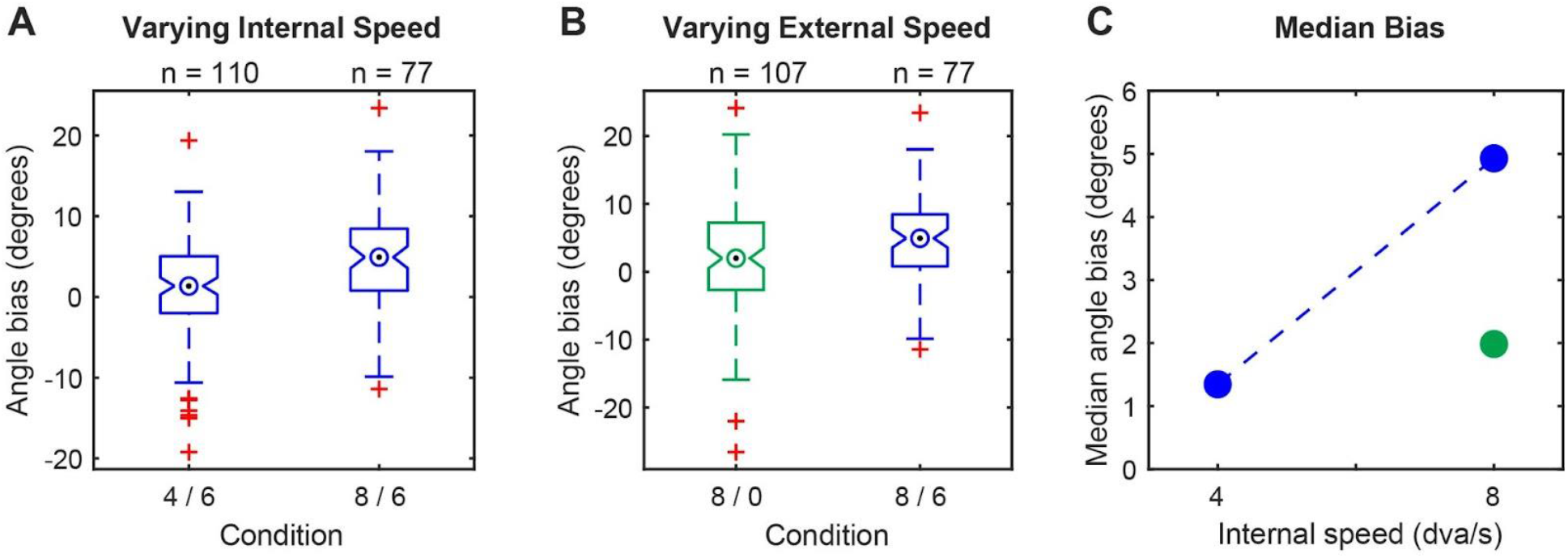
Saccade endpoints are parametrically biased by internal and external speed. A) Comparison of conditions with a fixed external speed of 6 dva/s and either 4 dva/s (labeled: 4 / 6) or 8 dva/s (labeled: 8 / 6) internal speed. B) Comparison of conditions with a fixed internal speed of 8 dva/s and either 0 dva/s (green; labeled: 8 / 0) or 6 dva/s (blue; labeled: 8 / 6) external speed. C) Summary of median angles for all three conditions. The blue dashed line shows the change observed by increasing only the internal speed. In panels, B and C, blue and green indicate external speeds of 6 and 0 dva/s, respectively.

To determine if the observed saccade bias is simply due to the internal motion as seen by Schafer and Moore (2007) in macaques, and not the combined effect of the internal and external drift, we used a Gabor stimulus with 8 dva/s internal speed and 0 dva/s external speed (Fig. 3B). In this condition, the saccade endpoints were slightly but significantly biased in the direction of internal motion (median angle bias = 1.99 deg.; p = 0.01, Wilcoxon signed rank test). However, the angle bias was substantially less than the matched condition with 8 dva/s internal motion and 6 dva/s external motion (1.99 deg. vs. 4.93 deg.; p = 0.01, Wilcoxon rank sum test).

Finally, we sought to determine if the amplitude or timing of saccades differed between conditions, which could potentially influence the observed saccade angles. Across all conditions and trial types, the mean saccade amplitude was 5.4 (SD = 1) dva and the mean saccade onset time was 239 (SD = 115) ms after the beginning of the saccade period. Between the conditions with a fixed external speed of 6 dva/s and a variable internal speed of 4 dva/s or 8 dva/s, we did not find any significant differences in the saccade amplitudes (p = 0.23, Wilcoxon rank sum test) or the saccade onset times (p = 0.97, Wilcoxon rank sum test). We also did not find any significant differences in the saccade amplitudes (p = 0.09, Wilcoxon rank sum test) or saccade onset times (p = 0.40, Wilcoxon rank sum test) between the conditions with a fixed internal speed of 8 dva/s and a variable external speed of 0 dva/s or 6 dva/s.

## Discussion

We trained two marmoset monkeys to report the location of a double drift stimulus with a saccade. Their saccade endpoints show a bias similar to the bias of human observers for saccades to targets on a radial path (Lisi & Cavanagh, 2022) and of human observers performing delayed saccades (Massendari et al., 2018), and the bias is modulated by changes in the external and internal velocity of the stimulus (Fig. 3C). Based on previous reports from humans (Kosovicheva et. al., 2014) and monkeys (Schafer & Moore, 2007), we expect that a static (no external drift) Gabor patch with fast internal motion will on its own bias saccade endpoints in the direction of internal motion. Indeed, we find a slight bias in the direction of internal motion for the static Gabor; however, when external motion is added the bias is more than doubled (Fig. 3C at 8 dva/s internal speed). These results demonstrate that the saccade targeting system of the marmoset monkey is biased by the double-drift illusion beyond the effect of internal motion alone. Remarkably, this occurs in the absence of any discernible differences in the amplitude or timing of saccades.

The saccade targeting system demonstrates a bias that is less than that seen for perceptual judgments. A simple explanation is that the oculomotor system does not integrate visual information over long timescales (Liu et. al., 2019; Lisi & Cavanagh, 2022). That is, it does not maintain and update a position estimate over extended periods of time, rather it uses only recent information to estimate the position of the stimulus. Figure 4 shows this pictorially. The black arrow indicates the physical position of the aperture over time. The dashed arrows indicate the position that is encoded in the perceptual (red) or oculomotor systems (blue). The perceptual system accumulates visual information over a long timescale, leading to a large perceptual offset. The oculomotor system, however, only integrates recent visual information and is subsequently less biased by the illusion. Temporal integration windows, which can be thought of as the capacity to maintain and update a representation over time, thus determine the magnitude of the effect.

**Figure 4:**
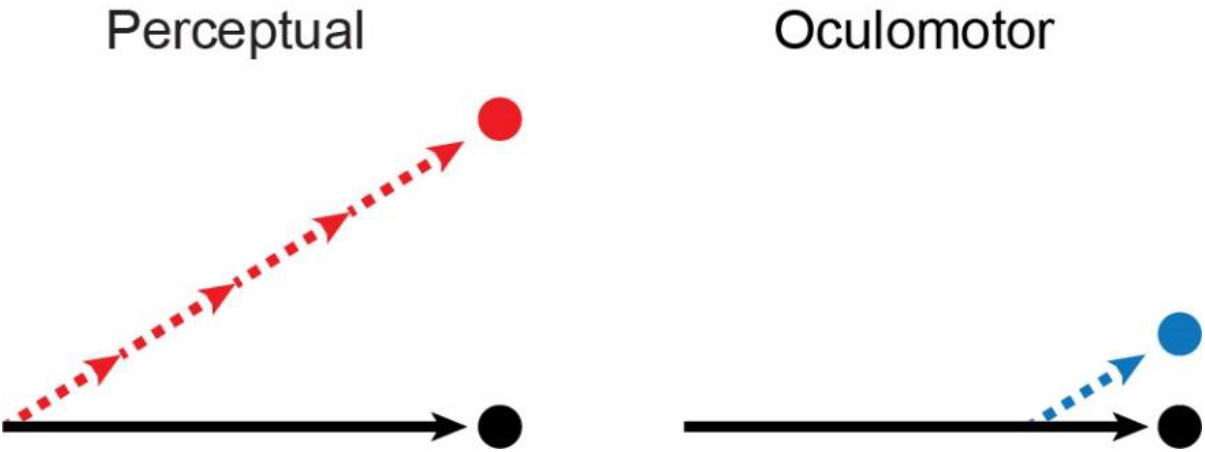
Illustration of the effect that different temporal integration times have on the position that is encoded by the perceptual and oculomotor systems. The black arrows represent the physical path of the aperture over time. The dashed arrows represent the stimulus paths that are encoded in the respective system. The perceptual representation (left) is depicted with a long integration time, while the oculomotor representation (right) is depicted with a short integration time. The colored circles on the right indicate the respective positions of the representations (physical = black; perceptual = red; oculomotor = blue).

The internal simulation that is constructed by our visual system is not a simple point-to-point mapping of the world. This is evidenced by visual illusions, which are capable of generating profound differences between the physical and the perceived, thus providing a unique tool for dissecting this process (Gregory, 1968; Eagleman, 2001). Non-human primates provide a crucial link for using visual illusions to study visual processing and perception at the level of single neurons (Sundberg et. al., 2006). Our results establish that the marmoset monkey oculomotor system is parametrically biased by the double-drift illusion, and suggest that marmosets may be experiencing effects similar to those seen by humans for the double-drift illusion. Many open questions remain about the sources of error, and the role of prediction (Palmer et. al., 2015). The sources of visual information utilized by the perceptual and oculomotor systems, and the degree to which they share visual information and neural substrates are also open matters of debate.

## References

Asaad, W. F., & Eskandar, E. N. (2008). A flexible software tool for temporally-precise behavioral control in Matlab. Journal of neuroscience methods, 174(2), 245–258.

Cavanagh, P., & Tse, P. U. (2019). The vector combination underlying the double-drift illusion is based on motion in world coordinates: Evidence from smooth pursuit. Journal of Vision, 19(14), 2–2.

Davis, Z. W., Muller, L., Martinez-Trujillo, J., Sejnowski, T., & Reynolds, J. H. (2020). Spontaneous travelling cortical waves gate perception in behaving primates. Nature, 587(7834), 432–436.

Davis, Z. W., Dotson, N. M., Franken, T., Muller, L., & Reynolds, J. (2022). Spike-phase coupling patterns reveal laminar identity in primate cortex. bioRxiv.

D’Souza, J. F., Price, N. S., & Hagan, M. A. (2021). Marmosets: a promising model for probing the neural mechanisms underlying complex visual networks such as the frontal–parietal network. Brain Structure and Function, 226(9), 3007–3022.

Eagleman, D. M. (2001). Visual illusions and neurobiology. Nature Reviews Neuroscience, 2(12), 920–926.

Gregory, R. L. (1968). Visual illusions. Scientific American, 219(5), 66–79.

Heller, N. H., Patel, N., Faustin, V. M., Cavanagh, P., & Tse, P. U. (2021). Effects of internal and external velocity on the perceived direction of the double-drift illusion. Journal of Vision, 21(8), 2–2.

Hwang, J., Mitz, A. R., & Murray, E. A. (2019). NIMH MonkeyLogic: Behavioral control and data acquisition in MATLAB. Journal of neuroscience methods, 323, 13–21.

Johnston, K., Ma, L., Schaeffer, L., & Everling, S. (2019). Alpha oscillations modulate preparatory activity in marmoset area 8Ad. Journal of Neuroscience, 39(10), 1855–1866.

Kosovicheva, A. A., Wolfe, B. A., & Whitney, D. (2014). Visual motion shifts saccade targets. Attention, Perception, & Psychophysics, 76(6), 1778–1788.

Lisi, M., & Cavanagh, P. (2015). Dissociation between the perceptual and saccadic localization of moving objects. Current Biology, 25(19), 2535–2540.

Lisi, M., & Cavanagh, P. (2022). Different integration of conflicting motion signals in perception and eye movements during object tracking. bioRxiv.

Liu, S., Yu, Q., Tse, P. U., & Cavanagh, P. (2019). Neural correlates of the conscious perception of visual location lie outside visual cortex. Current Biology, 29(23), 4036–4044.

Mitchell, J. F., Reynolds, J. H., & Miller, C. T. (2014). Active vision in marmosets: a model system for visual neuroscience. Journal of Neuroscience, 34(4), 1183–1194. PMCID: PMC3898283

Palmer, S. E., Marre, O., Berry, M. J., & Bialek, W. (2015). Predictive information in a sensory population. Proceedings of the National Academy of Sciences, 112(22), 6908–6913.

Schafer, R. J., & Moore, T. (2007). Attention governs action in the primate frontal eye field. Neuron, 56(3), 541–551.

Selvanayagam, J., Johnston, K. D., Schaeffer, D. J., Hayrynen, L. K., & Everling, S. (2019). Functional localization of the frontal eye fields in the common marmoset using microstimulation. Journal of Neuroscience, 39(46), 9197–9206.

Solomon, S. G., & Rosa, M. G. (2014). A simpler primate brain: the visual system of the marmoset monkey. Frontiers in neural circuits, 8, 96.

Steinberg, N. J., Roth, Z. N., Movshon, J. A., & Merriam, E. P. (2022). Neural Basis of The Double Drift Illusion. bioRxiv. 2022.01.25.477714; doi:https://doi.org/10.1101/2022.01.25.477714

Sundberg, K. A., Fallah, M., & Reynolds, J. H. (2006). A motion-dependent distortion of retinotopy in area V4. Neuron, 49(3), 447–457.

‘t Hart, B. M., Henriques, D. Y. P., & Cavanagh, P. (2022). Measuring the double-drift illusion and its resets with hand.

